# Diploidy alters the path of fluconazole adaptation in *Candida glabrata*

**DOI:** 10.64898/2026.05.22.727327

**Authors:** Benjamin Galeota-Sprung, Amy Fernandez, Cecilia Wright, Ronan Soto Tejada, Paul Sniegowski

## Abstract

*Candida glabrata* (syn. *Nakaseomyces glabratus*) is a major fungal pathogen, typically isolated as a haploid. We isolated a spontaneous diploid variant of the type strain CBS138 and performed experimental evolution under fluconazole. At physiologically relevant concentrations of fluconazole, we found that haploids and diploids both acquired *PDR1* mutations, as is commonly observed in *C. glabrata*; diploids additionally acquired heterozygous *ERG11* and *ERG25* mutations and were more likely to acquire aneuploidies. Despite an ancestral fitness advantage for haploids, after ∼200 generations the highest-fitness clones, as measured in competitive assays with and without fluconazole, were diploid double heterozygotes having nonsynonymous mutations to *PDR1* and *ERG11* or *PDR1* and *ERG25*. Diploid clones also had higher minimum inhibitory concentrations (MICs) to fluconazole. These results suggest, in this particular context, an evolvability advantage for diploids. In a follow-up experiment in which we rapidly increased the fluconazole concentration to a very high concentration, haploids and diploids adapted via entirely distinct routes: haploids via co-mutation in *ERG3* and *CgOSH3*, a previously unreported path to fluconazole resistance, and diploids via heterozygous mutation in *ERG25* coupled with trisomies of chrF, chrG, and chrI in all sampled clones, and chrC in most sampled clones.

**Importance:** *Candida glabrata* is a major human fungal pathogen that is typically, but not always, found as a haploid in clinical settings. The role of ploidy in the adaptation of *C. glabrata* to azole drugs had not previously been explored. We isolated a spontaneous diploid of the type strain CBS138 and compared its evolution, under physiologically relevant concentrations of fluconazole, to that of the haploid strain. After adaptation, the fittest and most resistant clones identified were diploids. In a second experiment, we evolved strains to very high concentrations of fluconazole: under this regime, haploids and diploids adapted by completely separate genetic pathways. These results suggest that ploidy can play an important role in the evolution of resistance of *C. glabrata* to clinically relevant drugs.

## Introduction

*Candida glabrata* (syn. *Nakaseomyces glabratus*) is a major fungal pathogen. In many parts of the world it is the second-leading cause of mucosal and invasive candidiasis, after *Candida albicans* [1], [2], [3], [4]. Its emergence as an important pathogen is relatively recent, as only in the last 30-35 years has *C. glabrata* become a major cause of candidiasis [5].

The recent reclassification of *C. glabrata* to *N. glabratus*, though both names remain in use, more accurately represents its phylogenetic placement: *C. glabrata* is only distantly related to *C. albicans* [6], [7]. It is instead closely related to other *Nakaseomyces* species, which together comprise a sister clade to *Saccharomyces*. Both *Saccharomyces* and *Nakaseomyces* share a common whole genome duplication in their evolutionary history.

While the predominant natural vegetative state of *Saccharomyces* species is diploidy, isolates of *C. glabrata* are usually found to be haploid, as is true for the majority of *Nakaseomyces* species. The *C. glabrata* genome includes the typical Saccharomycetaceae family triple cassette mating type system, but while there are both *MATa* and *MATalpha* lineages of *C. glabrata*, spontaneous *HO*-mediated mating-type switching apparently does not occur [8], [9]. Indeed, *C. glabrata* haploids have never been observed to mate. Nevertheless, genomic studies have provided compelling evidence for recombination in *C. glabrata* populations [6], [10], [11], suggesting that some mating does occur. While *MATa/MATalpha* diploids formed by syngamy have never been observed, some clinical isolates of *C. glabrata* have been found to contain a non-negligible minority of autodiploids [12]. Autodiploidization is a spontaneous doubling of genomic content that is common in *S. cerevisiae* [13].

Azole drugs are commonly used to treat fungal infections. Fluconazole (FLC) remains a first-line agent for mucosal candidiasis, though current global guidelines [14] emphasize susceptibility testing to guide treatment. For invasive candidiasis, echinocandins are now strongly recommended as first-line therapy across all *Candida* species because of their broad activity, safety profile, and challenges caused by azole resistance. *C. glabrata* is comparatively recalcitrant to FLC, compared to other fungal pathogens, in two aspects. First, *C. glabrata* is naturally less susceptible to FLC than *C. albicans* by an order of magnitude as quantified by minimum inhibitory concentration (MIC). Second, *C. glabrata* easily acquires mutations to *PDR1* that drive resistance to fluconazole [15], [16], [17], [18]. *CgPDR1* is the master transcriptional regulator of three separate ABC transporter efflux pumps (Cdr1, Pdh1, and Snq2). The predominance of *PDR1* gain-of-function mutations as the primary azole resistance mechanism in *C. glabrata* is unique among the major fungal pathogens and is a function of its phylogeny, as the CUG-Ser clade fungal pathogens *C. albicans, C. tropicalis, C. parapsilosis, and C. auris* share a different transcriptional architecture.

Azole drugs such as fluconazole inhibit the 14α-demethylase CYP51 (Erg11), blocking the C-14 demethylation of lanosterol, a key step in ergosterol biosynthesis. Ergosterol is a key membrane component in fungi, analogous to cholesterol in mammals. While the resulting depletion in ergosterol is a difficulty for the cell, the fungistatic consequence of Erg11 inhibition is thought to be primarily driven by accumulation of 14α-methylated sterol intermediates that Erg3 processes into the “toxic diol” 14α-methylergosta-8,24(28)-dien-3β,6α-diol [19], [20].

Many evolution experiments conducted in yeast species have investigated the role of ploidy in adaptation [21], [22], [23], [24], [25], [26]. On balance, it seems that these experiments tend to find an adaptive advantage for haploids over diploids, but whether haploids or diploids (or higher ploidies) have an easier adaptive path in any scenario is contingent upon the particulars. For example, Anderson et al (2004) [27] found that at low concentrations of fluconazole, *S. cerevisiae* diploids adapted faster via partially-dominant *PDR1* mutations (due to increased mutational target), while at high concentrations haploids adapted faster via loss-of-function (LOF) mutations to *ERG3*. Considering the rate of adaptation in the abstract, diploids face some disadvantages compared to haploids. Most obviously, fully recessive beneficial mutations are invisible to natural selection [28] unless and until some process (meiosis followed by syngamy, or various loss-of-heterozygosity mechanisms) creates homozygosity. Conversely, diploids have the advantage of twice the mutational target of haploids, as well as the possibility of higher-fitness heterozygous states (heterozygote advantage, or overdominance) that are inaccessible to haploids. Diploids also experience large-scale mutations such as copy number variation in chromosome number (aneuploidy) at a higher rate than haploids [29], and can tolerate gain-of-copy-number aneuploidies more easily, the relative change in dosage from disomy to trisomy being smaller than the change from monosomy to disomy. In *C. albicans*, the segmental aneuploidy i(5L) confers FLC resistance [30]. Tolerance and adaptive potential of aneuploid states may scale with ploidy, in general [24].

The consequences of ploidy for experimental evolution of antifungal resistance in *C. glabrata* have not previously been investigated. Considering the clinical importance of azole resistance, and broad evolutionary interest in questions of ploidy, we were motivated to conduct experiments comparing the adaptation of haploid and diploid strains of *C. glabrata* to fluconazole. To our knowledge these are the first evolution experiments reported for diploids of this species. In our primary experiment, we held the concentration of fluconazole static at 50 or 100 μg/mL, for ∼200 generations. We chose these conditions to reflect the highest steady state concentration in plasma likely to be seen in clinical practice [31], [32], [33], [34]. We found that the highest-MIC and highest-fitness (as measured by competition assays in both YPD and YPD + FLC) clones emerging from this evolution experiment were diploid. In a follow-up experiment conducted over a shorter time period, we increased the concentration of fluconazole dynamically as the populations evolved, to concentrations well above clinical exposure. At these very high concentrations, adaptation across ploidies revealed no shared mutational targets between haploids and diploids.

## Methods

### Strains and growth conditions

The haploid MATalpha *C. glabrata* strain CBS138 (ATCC 2001) was the ancestor for all work described here. We refer to this strain as H-anc. A spontaneous autodiploid clone of this strain was isolated as described below, and cataloged as YPS3723; we refer to this strain as D-anc. Growth conditions for all experiments and assays, except where noted otherwise, were as follows: 10 mL liquid volume in 50 mL Erlenmeyer-type glass flasks, at 37° C with shaking at 200 rpm, in Yeast Peptone Dextrose (YPD; 1% yeast extract, 2% peptone, and 2% glucose) medium sometimes supplemented with fluconazole.

### Isolation of spontaneous diploids

Overnight cultures of CBS138 grown as described above were diluted and spread onto YPD agar plates containing 5 μg/mL phloxine B (Acros Organics 189470050) at ∼140 colonies per plate. Plates were grown at 37°C for 24 hours after which colonies were allowed to grow slowly at room temperature. Spontaneous diploids appeared as colonies with a darker shade of pink to red than the wild-type [12].

### Determination of ploidy via flow cytometry

Ploidy was assayed by flow cytometry using a modified version of the protocol described in [35]. Overnight cultures were sub-cultured until mid-log phase (OD600 ∼ 0.8). 1 mL aliquots were spun down and resuspended in 70% ethanol and incubated at RT for 1 hour or overnight at 4 C. 500 uL aliquots were then washed once in 50 mM sodium citrate buffer, resuspended in the same buffer, and sonicated using a Fisher Scientific/Branson Sonifier Model 150 Cell Disruptor for two 5 s pulses at 40% power separated by 8 s rest. Samples were then spun down and resuspended in 50 mM sodium citrate buffer + 0.5 mg/mL RNase A (Roche 10109169001) and incubated at 37 C for 2.5 h, after which samples were spun down and resuspended in 500 uL 50 mM sodium citrate buffer + 2 μg/mL propidium iodide (Thermo P3566) and incubated at 37 C overnight. Samples were diluted using the same buffer + propidium iodide to a density of ∼1.5e05 cells/mL and loaded onto a 96-well plate for analysis using a Guava EasyCyte HT benchtop flow cytometer outfitted with a blue (488 nm) laser. Ploidy was determined by observing the location of the first and second peaks on the RED channel. In some instances, a FACSCalibur (Becton Dickinson) flow cytometer was used instead, for which 10 μg/mL propidium iodide was used with all other preparatory steps the same. Signal height on the red (PI) channel was analyzed in R.

### Adaptation to fluconazole at fixed concentration

Beginning from single colonies, we inoculated 3 replicate populations of H-anc and D-anc in 6 mL YPD, which were subsequently transferred to YPD, YPD + 50 μg/mL FLC, and YPD + 100 μg/mL FLC, for a total of 2 x 3 x 3 = 18 populations. Throughout, population names such as H50-A indicate founding ploidy (H or D), FLC concentration (0, 50, or 100 μg/mL), and replicate (A–C). These populations were propagated for ∼200 generations by transferring 10 uL into 10 mL fresh medium for 20 transfers. Transfers were performed every ∼72 h as pilot experiments showed that initial growth in YPD + FLC would be slow. At regular intervals, 100 uL aliquots of populations were mixed 1:1 with 30% glycerol and stored at -80 C. Populations were regularly plated to YPD agar supplemented with 5 μg/mL phloxine B in order to assess ploidy. One of the three replicates of the diploid-founded populations at 100 μg/mL FLC was lost to technical error. Up to two clones from each population’s final transfer were isolated, grown up, and stored for sequencing. These clones were randomly chosen from the major lineages present as estimated by colony color. Clones isolated in this way are referred to by a terminal number appended to the population name. For example, D100-A1 and D100-A2 are the two clones isolated from the final growth cycle of the A replicate of the diploid-founded 100 μg/mL FLC condition.

### Adaptation to dynamic concentration of fluconazole

In this experiment, 4 replicate populations of H-anc and D-anc were transferred into a range of concentrations of FLC, in a volume of 10 mL. The flask with good growth (OD600 > 1.2) and the highest FLC concentration was chosen as the source for the next transfer. Transfers occurred every 48 h and continued for 5 growth cycles. To standardize the number of cells transferred, the transfer volume was 100 uL for H-anc and 200 uL for D-anc. One representative clone was randomly chosen from each population at the end of the experiment for sequencing. Clones isolated from this experiment are referred to by catalog number (e.g. YPS4228).

### DNA extraction and whole-genome sequencing

The annotated CBS138 genome version s05-m03-r47 (www.candidagenome.org) was used as the reference for all genetic analyses. For each sequenced clone, after growth overnight in YPD, DNA was extracted from 1 mL of culture using kit D7005 from Zymo Research. Short-read whole-genome sequencing was performed on an Illumina NovaSeq X Plus sequencer in multiplexed shared-flow-cell runs, producing 2 × 151 bp paired-end reads. The median of medians read depth across all samples was 29x. An initial ploidy of 1 or 2 as determined by flow cytometry was assigned to each clone. Variants were called as follows. Paired-end Illumina reads (pre-trimmed by the sequencing facility) were QC-profiled with fastp v1.3.2 in report-only mode and aligned to the reference genome with bwa-mem2 v2.3. Alignments were coordinate-sorted, fixmate-tagged and PCR duplicates were marked with samtools v1.23.1. Per-chromosome copy number was estimated from 5 kb-binned coverage with mosdepth v0.3.13; integer ploidy assignments per chromosome, as well as partial-arm events ≥100 kb after masking subtelomeres and known repetitive loci, were written to per-sample CNV-map BED files. Nuclear variants were jointly called with freebayes v1.3.10 in two cohorts, one comprising the haploid ancestor and its descendants, the other the diploid ancestor and its descendants, with each sample’s per-region copy number supplied via --cnv-map to handle aneuploidies. The mitochondrion was called separately with freebayes --ploidy 1 --pooled-continuous to obtain per-site allele frequencies for the high-copy, potentially heteroplasmic mtDNA. Variants present in the matched ancestor were subtracted from each evolved sample. Sites with less than 8x ancestral read depth were removed from further analysis; this comprised 0.04% and 0.03% of the genome for the haploid and diploid ancestors, respectively. Surviving variants were annotated with SnpEff v5.4.0c against a custom SnpEff annotation index built from the matching GFF3, retaining HIGH, MODERATE, and LOW coding effects plus regulatory MODIFIER variants within 1 kb upstream of a TSS and in annotated UTRs. Results were spot-checked with breseq [36].

### Phasing of *PDR1* mutations

Clones heterozygous for two *PDR1* mutations (D50-A2 and D100-C1) were phased as follows. Primers 5’-GCCCACACTTGCTTGGTTTT-3’ and 5’-ACGAAGGTAGTGCTGAGGTT-3’ were used to amplify a 3984 bp region that included the entire *PDR1* gene. Amplicons were cleaned up, pooled, and sequenced using the ONT GridION with an R10.4.1 flow cell in 400 bps mode (adaptive sampling disabled). The run yielded 2,663,498 reads (mean length 3,697 bp; 93.6% at Q ≥ 9), of which ∼1.9 million were full length and retained for the analysis. Reads were mapped to the CBS138 genome using minimap2 v2.30. Per-read genotypes at the four variant positions were extracted from samtools mpileup with the --output-QNAME option. Reads were classified as supporting mutations in *trans* if they contained one mutation and *cis* if they contained two mutations.

### Fitness competitions

Head-to-head (two-way) haploid vs. diploid fitness competitions were conducted by making use of the colony color difference between diploid and haploid strains. Clones were competed in the condition in which they evolved, either YPD + 50 μg/mL FLC or YPD + 100 μg/mL FLC. Only clones that evolved in the same condition were competed together. Day 1: clones were grown overnight in YPD + FLC. Day 2: for each competitor, 10 uL of culture was transferred into a fresh 10 mL YPD + FLC. Day 3: After 24 h growth, 4 replicate competitions in 4 separate flasks were set up by transferring 3.3 uL haploid competitor and 6.7 uL diploid competitor into 10 mL fresh YPD + FLC, in order to start with ∼50% frequency of each competitor. An aliquot from each flask was then further diluted and spread onto YPD + phloxine B agar plates to assay the initial frequency of the competitors. Competition cultures were grown for 72 h at 37 C, replicating the conditions of the evolution experiment, and then aliquots from each replicate diluted and spread onto YPD + phloxine B agar plates to assay the final frequency of the competitors. Selection coefficients were computed as the time-averaged difference in Malthusian parameter from the frequency data according to the formula

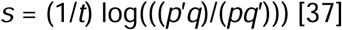

where *p* is the initial frequency of one competitor, *q=1-p* is the initial frequency of the other competitor, and the final frequencies are *p*′ and *q*′*=1-p*′ respectively; *t* is the number of generations, assumed to be log_2_(1000)=9.97. This formula can equivalently be written

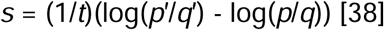

For each head-to-head competition, the *p*-value that the mean *s* across the 4 replicates was significantly different from zero was computed via two-sided t-test.

Multi-strain (five-way) competitions were conducted using each strain’s *PDR1* mutation as a barcode; competitors were chosen to each have a unique *PDR1* mutation. Competitions were performed in both YPD and YPD + 100 μg/mL FLC, with two replicates for each condition. Competitor strains were grown up in 6 mL YPD overnight and then transferred to both YPD and YPD + 100 μg/mL FLC flasks, and grown for 48 h. To initiate the competition, 1 mL of culture of each competitor was combined. From this pre-competition mix, 1 mL was used for DNA extraction, and 10 uL was used to initiate the competition in fresh medium. Competition cultures were grown for two 48-hour cycles, at which point another 1 mL culture was used for DNA extraction.

From the initial and final DNA extractions for each replicate, the same primers described above were used to amplify a region that included the entire *PDR1* gene. Amplicons were cleaned up and sequenced using the Illumina platform as described above. Initial and final frequencies of each competitor were computed using breseq in polymorphism mode [36], with the amplicon as the reference sequence. Median coverage was approximately 10^5^ reads/site. Selection coefficients were computed from the frequency data according to the formula above, with *t* = 2log_2_(1000) = 19.93 generations, choosing the lowest-fitness competitor to be the reference against which selection coefficients were computed for the others. Statistical tests of whether the difference in *s* between two strains differed from zero were conducted by first fitting a linear model with competition replicate and strain as fixed effects. The model was fitted separately for each environment. Then, each diploid was compared with each haploid using two-sided t-tests on linear contrasts from the fitted model. Holm’s correction, applied jointly across all comparisons, adjusted the resulting *p*-values.

### MIC assays

Minimum inhibitory concentration (MIC) assays were conducted in 96-well plates. Clones were started from single colonies and grown overnight in 6 mL YPD, from which 100 uL was transferred to 10 mL fresh YPD. After 24 h growth, a 1/100 dilution in fresh YPD was prepared and 10 uL of the resulting dilution was added to wells of a 96-well plate containing 190 uL of YPD + FLC such that the final concentrations spanned the powers of two from 16 μg/mL to 2048 μg/mL FLC. Plates were grown for 20 h at 37 C without shaking. Wells were pipetted up and down to mix and then read at 600 nm on a BioTek Elx808 plate reader. Two columns were reserved as blanks. The cutoff for scoring a well for growth was 0.10 OD600 units after subtracting the mean blank value from all wells; this value corresponds to a ∼93% inhibition of final OD600 in the ancestors.

## Results

### Isolation of spontaneous diploids

We conducted a screen of CBS138 (H-anc) for diploids, scoring colonies from populations grown in YPD and spread on YPD + phloxine B agar plates. After growth, diploid colonies are a darker pink-to-red color than are haploid colonies [12]. The mechanism behind this color difference is not currently established. We observed that petite colonies lacking mitochondrial function are darker still, in addition to being much smaller; we did not choose these. Out of ∼8000 colonies examined, we isolated 10 potential diploid colonies based on color and size. Flow cytometry confirmed that 9 were diploid, a frequency of ∼0.1%. One diploid isolate, cataloged as YPS3723 (D-anc; Fig. 1A), was the ancestral strain for the subsequent experiments described here. Whole-genome sequencing of this strain revealed no heterozygosities, including in the *MAT* region, supporting that it is a MATα/MATα autodiploid (Fig. S1A). We found that at saturation in YPD, diploid culture density was about 55% of haploid culture density (Fig. S2).

**Fig. 1.**
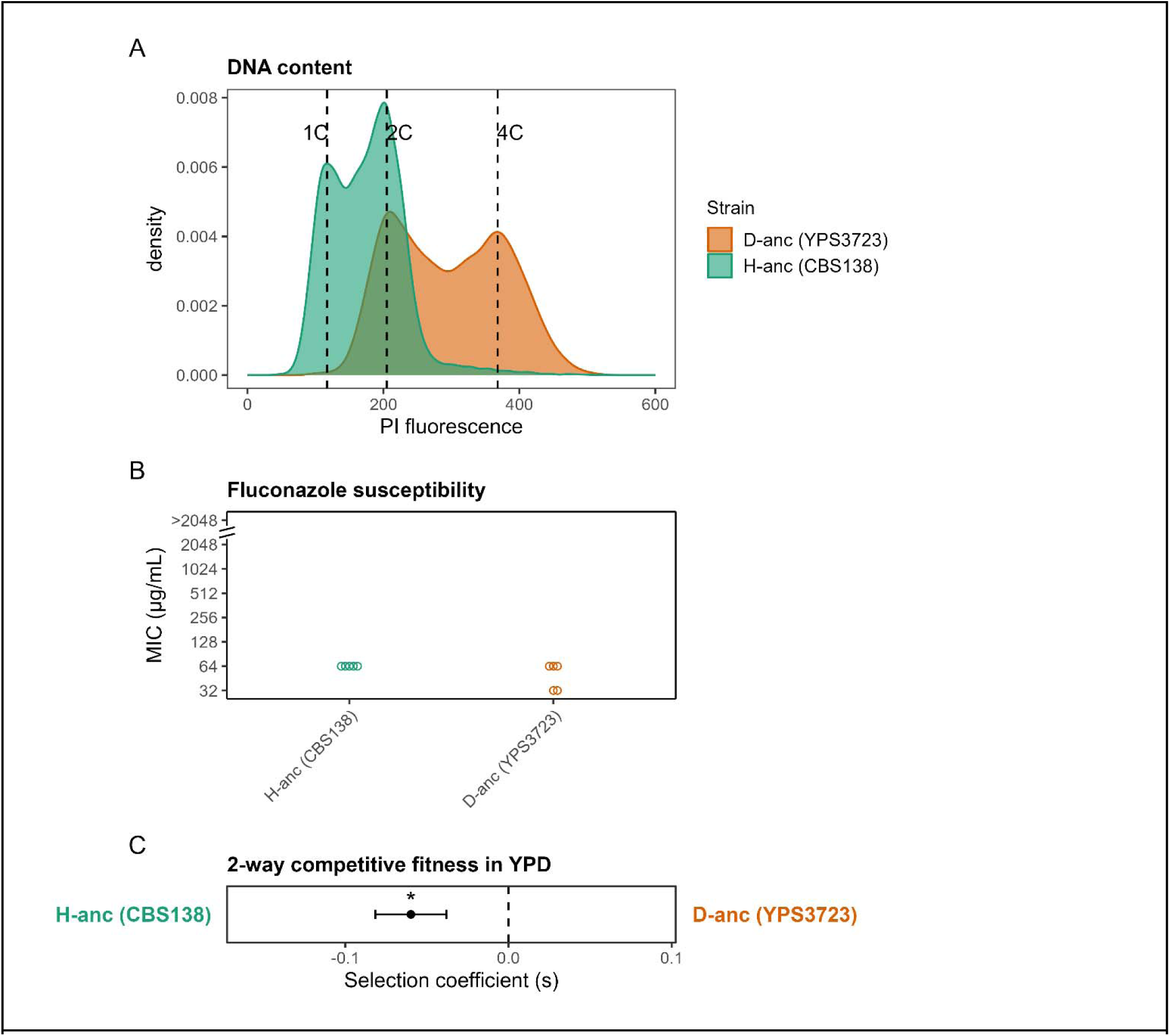
(A) The diploid ancestor (D-anc) is a spontaneous diploid isolate of the haploid ancestor (H-anc; CBS138). (B) Fluconazole MICs of both ancestors. (C) In a competitive fitness assay conducted in YPD, the haploid ancestor is fitter than the diploid ancestor (*p* = 0.012).

MIC tests conducted in YPD revealed a slight difference in fluconazole sensitivity between the two ploidies. We measured the MIC using a twofold dilution series in YPD (Fig. 1B). For H-anc, we measured an MIC of 64 μg/mL across all replicates. For D-anc, the MIC was variously measured at either 64 or 32 μg/mL across replicates, reflecting a pattern of consistently intermediate growth at 32 μg/mL.

We competed H-anc and D-anc in head to head fitness competitions in which the relative frequency of the two ploidies over time was estimated by exploiting the colony color difference when grown on phloxine B. We found the haploid to be substantially more fit than the autodiploid in YPD (Fig. 1C); in contrast, this is not the case for *S. cerevisiae* [39], [40]. Interestingly, we observed that the fitness difference between haploid and diploid *C. glabrata* is abolished in minimal media (Fig. S3).

### Adaptation to a fixed concentration of fluconazole

We conducted an evolution experiment with 18 populations in total: two ancestral strains (the haploid CBS138 and the diploid YPS3723, referred to as H-anc and D-anc respectively) ✕ three environments (YPD, YPD + 50 μg/mL FLC, and YPD + 100 μg/mL FLC) ✕ threefold replication of each strain-environment combination. Pilot experiments conducted with small populations cultured in 96-well plates, transferred daily, tended to enrich for mitochondrial dysfunction (petite mutants), a known resistance mechanism [41]. Therefore we chose to propagate experimental populations in flasks, with longer intervals between transfers. Relative to the pilot, this design increased the mutational supply via larger population size, and permitted additional mutational input from any subpopulations capable of slow growth above the MIC (tolerance) [42]. Populations were propagated for 20 transfers with a dilution factor of 1000 at each transfer, resulting in ∼200 generations.

Populations were monitored for shifts in ploidy by regular plating to YPD agar + phloxine B. Light-colored lineages arising in diploid populations were sometimes but not always haploid, as determined by flow cytometry. One diploid population (D100-B) was lost due to technical error. We detected no ploidy shifts in the no-FLC condition, nor in any haploid-founded population, while two of three diploid-founded populations at 50 μg/mL FLC were majority-haploid by final transfer (Fig. 2A). At 100 μg/mL FLC, all of the diploid-founded populations remained diploid, though we detected a haploid lineage in replicate A that attained a maximum frequency of ∼25% at the 12th transfer and was not detectable by the final passage.

**Figure 2.**
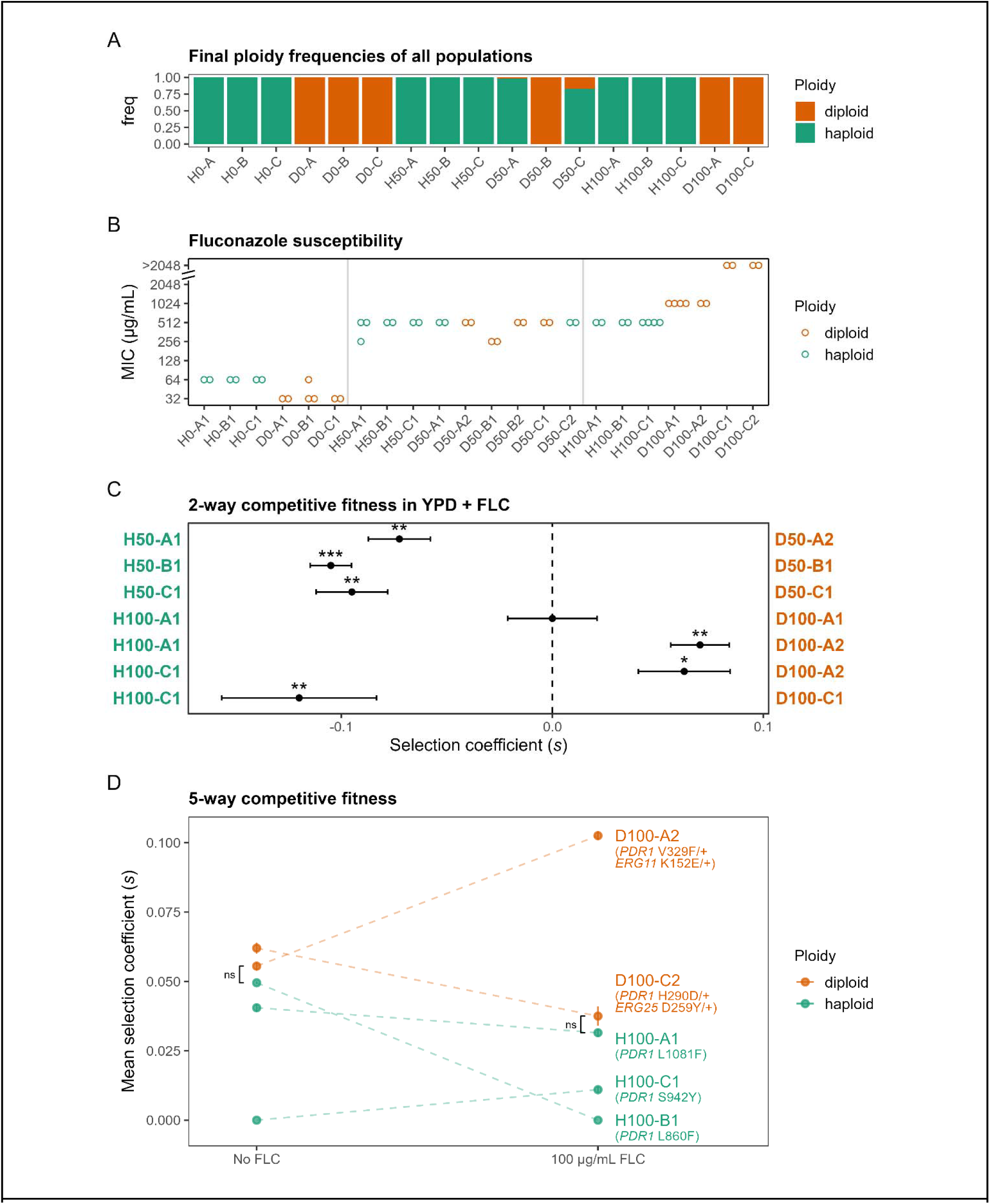
(A) The final ploidy frequencies of the evolved populations; note that D50-A and D50-C were diploid-founded. (B) MICs of clones isolated from populations evolved in 0, 50, and 100 μg/mL FLC. (C) Head-to-head fitness competitions of evolved haploid and diploid clones, performed in YPD with 50 or 100 μg/mL of FLC, according to the evolution condition. Stars indicate that the mean *s* in a 2-way competition differs significantly from 0: ∗, *p*< 0.05; ∗∗, *p*< 0.01; ∗∗∗, *p*<0.001. (D) 5-way fitness competitions among clones evolved in 100 μg/mL FLC, performed in both YPD and YPD + 100 μg/mL FLC. Within each environment, all haploid to diploid differences in mean s between two clones differed significantly from zero (*p* < 0.01) except for those indicated by “ns” (see Methods for details of statistical procedure). Dotted lines connect a single clone’s measured selection coefficient in the two environments. The genotypes of the clones are indicated.

We sequenced the genomes of two clones from the final growth cycle of each diploid-founded population and of one clone from the final transfer of each haploid-founded population. All sequenced haploids possessed the same V713I substitution to CAGL0D05280g (the ortholog of *MET10*); this was presumably a founder effect and we did not find any reports of an association between *MET10* and azole resistance. All FLC-adapted clones from the final growth cycle had nonsynonymous *PDR1* mutations including N283D, G558V, and D954N (Table 1). Haploid clones isolated from diploid-founded populations shared the same *PDR1* mutation as surviving diploids (compare the genotypes of D50-A1 and D50-A2, and D50-C1 and D50-C2), indicating that the mutation originally arose in a diploid background that subsequently founded a haploid lineage. With respect to *PDR1* mutations, diploid clones were variously (1) heterozygous for a single mutation; (2) homozygous for a single mutation; or (3) heteroallelic for two separate *PDR1* mutations (Fig. S4). In all, we observed 11 distinct *PDR1* mutations. One *PDR1* mutation, L860F, was observed in three independent replicates. The mutations showed similar clustering into three regions as observed in Ferrari et al. (2009) [17].

**Table 1.** Ploidies, genotypes, and karyotypes of clones that evolved over ∼200 generations (∼100 generations in the case of the T10 clones) at 50 or 100 μg/mL FLC, as determined by whole-genome sequencing.

| **Clone** | **Transfer** | **Catalog** | **Ancestor** | **Ploidy** | **Genotype** |
|---|---|---|---|---|---|
| H50-A1 | 20 | YPS4119 | haploid | haploid | *PDR1* G583D |
| H50-B1 | 20 | YPS4120 | haploid | haploid | *PDR1* L860F |
| H50-C1 | 20 | YPS4121 | haploid | haploid | *PDR1* L860F |
| H100-A1 | 20 | YPS4122 | haploid | haploid | *PDR1* L1081F |
| H100-B1 | 20 | YPS4123 | haploid | haploid | *PDR1* L860F |
| H100-C1 | 20 | YPS4124 | haploid | haploid | *PDR1* S942Y |
| D50-A1 | 20 | YPS4146 | diploid | *haploid* | *PDR1* D954N |
| D50-A2 | 20 | YPS4147 | diploid | diploid | *PDR1* D954N/D954N |
| D50-B1 | 20 | YPS4148 | diploid | diploid | *PDR1* N283D/E579G |
| D50-B2 | 20 | YPS4149 | diploid | diploid | *PDR1* N283D/+; chrE[335 kb+]x3 chrI[715kb+]x3 |
| D50-C1 | 20 | YPS4150 | diploid | diploid | *PDR1* G558V/G558V |
| D50-C2 | 20 | YPS4151 | diploid | *haploid* | *PDR1* G558V |
| D100-A1 | 20 | YPS4125 | diploid | diploid | *PDR1* V329F/+ *ERG11* K152E/+; chrCx3 |
| D100-A2 | 20 | YPS4126 | diploid | diploid | *PDR1* V329F/+ *ERG11* K152E/+ |
| D100-A3(T10) | 10 | YPS187F03 | diploid | diploid | *PDR1* V329F/+ *ERG11* K152E/+; chrCx3 chrFx3 chrIx3 |
| D100-A4(T10) | 10 | YPS187F05 | diploid | *haploid* | *ERG11* K152E/+; chrEx2 |
| D100-C1 | 20 | YPS4154 | diploid | diploid | *PDR1* V329F/L935F *ERG25* D259Y/+ |
| D100-C2 | 20 | YPS4155 | diploid | diploid | *PDR1* H290D/+ *ERG25* D259Y/+ |

Diploid clones evolved at 100 μg/mL FLC had nonsynonymous mutations to one or the other of two additional putative resistance-associated loci, *ERG11* and *ERG25*. *ERG11* encodes lanosterol 14α-demethylase, the target of fluconazole, while *ERG25* encodes a C-4 sterol methyl oxidase that catalyzes the first oxidative step in removal of the two C-4 methyl groups from sterol intermediates in the ergosterol biosynthesis pathway. Two final diploid clones were found to have chromosomal aneuploidies: D50-B2 carried partial trisomies of ChrE (including *ERG11*) and ChrI, while D100-A1 carried a complete trisomy of ChrC.

We conducted MIC assays on all clones isolated from the final growth cycle (Fig. 2B). MICs for clones evolved under FLC generally increased by about an order of magnitude. The diploid clones with *ERG11* or *ERG25* mutations had higher MICs than clones possessing only *PDR1* mutations.

As mentioned above, one diploid-founded population evolving at 100 μg/mL FLC (D100-A) experienced invasion by a spontaneous haploid lineage that rose to intermediate frequency partway through the experiment, but was not detectable by the final growth cycle. We additionally sequenced one haploid and one diploid clone from the 10th transfer (∼100 generations) of this population. The intermediate haploid clone was lacking a *PDR1* mutation and possessed a disomy of chrE that allowed maintenance of the same heterozygous *ERG11* K152E mutation detected in diploids from this population. The intermediate diploid clone possessed the same *PDR1* V329F and *ERG11* K152E heterozygous mutations as the final clones from this population, shared a chrC trisomy with one final clone, and also had additional trisomies to chrF and chrI not present in either final clone.

### Haploid *vs.* diploid fitness competitions

We asked whether, in the environment in which they evolved, haploid or diploid clones were more fit. We first investigated this by conducting head-to-head fitness competitions (Fig. 2C) in YPD + 50 or 100 μg/mL FLC. We could not use the ancestors in these conditions, due to their very substantial fitness deficit under fluconazole compared to the evolved clones, so we competed evolved haploid and diploid clones against each other, pairing clones from the A, B, and C replicates of each environment. In these assays, the relative frequency of two competitors was estimated, using the colony color difference when plated on YPD + phloxine B, at the start and end of one growth cycle. Because some lineages evolved a lighter colony color, we could not compete all diploid clones in this assay. For example, D100-C2 proved too difficult to discriminate from haploids based on color. Consistent with their ancestral growth advantage, haploids were often fitter than diploids in these assays. Remarkably, though, the evolved diploid D100-A2 with genotype *PDR1* V329F/+ *ERG11* K152E/+ was substantially and significantly fitter than the evolved haploid clone it was tested against. To extend this finding, we conducted an extra crosswise head-to-head fitness competition with this clone, matching to H100-C1, against which it displayed a similar fitness advantage.

We conducted additional five-way fitness competitions among clones evolved in 100 μg/mL FLC, via a different method. Competitors were chosen so that a unique *PDR1* mutation served as a barcode for sequencing-based frequency estimation. These competitions were carried out both with and without FLC. These experiments confirmed our previous finding that D100-A2 (*PDR1 ERG11* double heterozygote) was significantly fitter than all representative evolved haploid clones in YPD + 100 μg/mL FLC. The other diploid tested in this assay, D100-C2 (*PDR1 ERG25* double heterozygote), was significantly fitter than two of three haploid competitors in this condition. In YPD without FLC, D100-A2 was significantly fitter than two of three haploid competitors, and D100-C2 was significantly fitter than all three haploid competitors. Thus, in the 100 μg/mL FLC evolution condition, some evolved diploid clones were unconditionally fitter (*i.e.* fitter in both YPD and YPD + FLC) than evolved haploid clones.

### Adaptation under dynamically increasing fluconazole concentrations

The genetics of adaptation in the primary evolution experiment suggested that at higher concentrations of FLC, diploids explore different adaptive pathways than do haploids. We therefore conducted an additional evolution experiment, allowing populations to adapt to a concentration of FLC that was dynamically adjusted upwards to a limit of 1024 μg/mL. Although such concentrations exceed those achievable in candidiasis treatment, we reasoned that high concentrations may amplify adaptive differences across the two ploidies.

Across all populations, adaptation to high concentrations of FLC was rapid (Fig. 3A). We detected no whole-genome ploidy shifts in this experiment. We sequenced one representative clone from each replicate population, and the resulting mutation sets displayed a high degree of parallelism within each ploidy (Table 2). All independently isolated haploid clones carried mutations to *ERG3* and CAGL0J10780g, many clearly loss-of-function. The latter locus is the ortholog of *ScOSH3* and we henceforth refer to it as *OSH3*. All four diploid clones carried a heterozygous mutation to *ERG25*, two clearly loss-of-function. All diploid clones carried trisomies of chrF, chrG, and chrI; three of four also carried a trisomy of chrC. Thus, in this experiment, haploids and diploids adapted via distinct routes: haploids fixed mutations in genes affecting sterol biosynthesis and trafficking (*ERG3* and *OSH3*), while diploids combined heterozygous mutations in a different sterol biosynthesis gene (*ERG25*) with large-scale aneuploidy.

**Figure 3.**
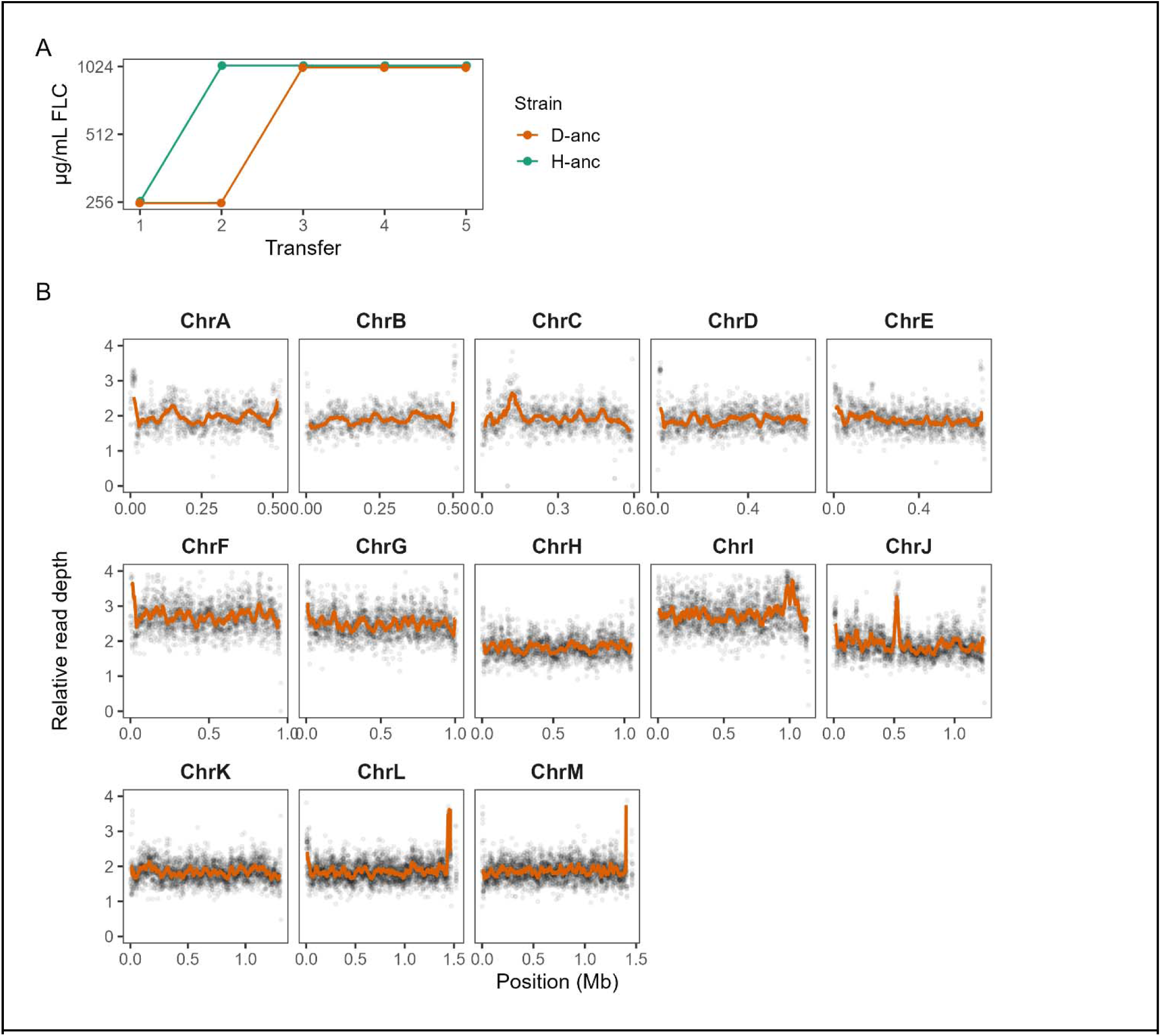
(A) The concentration of FLC was adjusted dynamically upwards to 1024 μg/mL. All 4 replicates of each ploidy ollowed the same concentration trajectory. (B) Relative read depth, by chromosome, of one representative clone YPS4254) isolated from a diploid-founded replicate, displaying trisomies of ChrF, ChrG, and ChrI.

**Table 2.** The haploid and diploid genotypes that evolved under adaptation to increasing concentrations of FLC.

| Transfer | Rep | Catalog | Ancestor | Ploidy | Genotype |
|---|---|---|---|---|---|
| 5 | 1 | YPS4226 | haploid | haploid | *ERG3* \*365Y *OSH3* R643I |
| 5 | 2 | YPS4228 | haploid | haploid | *ERG3* \*365Y *OSH3* D936N |
| 5 | 3 | YPS4230 | haploid | haploid | *ERG3* T77fs *OSH3* D930E |
| 5 | 4 | YPS4232 | haploid | haploid | *ERG3* H293Y *OSH3* G457fs |
| 5 | 1 | YPS4254 | diploid | diploid | *ERG25* D259H/+; chrF x3 chrG x3 chrI x3 |
| 5 | 2 | YPS4256 | diploid | diploid | *ERG25* E113\*/+; chrC x3 chrF x3 chrG x3 chrI x3 |
| 5 | 3 | YPS4258 | diploid | diploid | *ERG25* C70F/+; chrC x3 chrF x3 chrG x3 chrI x3 |
| 5 | 4 | YPS4260 | diploid | diploid | *ERG25* T226fs/+; chrC x3 chrF x3 chrG x3 chrI x3 |

## Discussion

### The fittest clones after adaptation to 100 μg/mL FLC are diploid

In this work, we investigated the influence of ploidy on the evolution of resistance to fluconazole in *C. glabrata*. To do so, we first isolated a spontaneous autodiploid from CBS138. After ∼200 generations of adaptation to 50 or 100 μg/mL FLC, haploids uniformly adapted via single *PDR1* mutations. Diploids also acquired *PDR1* mutations that were variously homozygous, heterozygous, or heteroallelic. Diploids adapted to 100 μg/mL FLC had additional mutations to *ERG11* or *ERG25*, and we also detected aneuploidies in end-of-experiment diploids but not haploids. These diploid clones had higher MICs to FLC than did haploid clones (Fig. 2B). Remarkably, given that diploids began the experiment at a substantial fitness disadvantage to haploids, the evolved diploids that are *PDR1 ERG11* or *PDR1 ERG25* double heterozygotes are generally fitter than evolved haploid clones, increasing in frequency in competition in both YPD alone and YPD + 100 μg/mL FLC (Figs. 2C, D). In particular, the *PDR1 ERG11* double heterozygote is exceptionally fitter than any tested haploid in YPD + 100 μg/mL FLC.

Although the fittest clones after adaptation to 100 μg/mL were diploid, at 50 μg/mL FLC diploid populations were vulnerable to invasion by lineages that had undergone spontaneous reversions to haploidy (Fig. 2A). At 100 μg/mL FLC, we detected an ultimately unsuccessful spontaneous haploid lineage partway through the experiment. All sequenced diploids (Table 1) carried at least two putatively adaptive mutations: either homozygous/heteroallelic mutations to *PDR1*, or a single heterozygous mutation to *PDR1* plus an additional mutation to an *ERG* gene and/or an aneuploidy. We interpret these observations as follows. A baseline fitness disadvantage for diploids, combined with a non-negligible rate of reversion from diploidy to haploidy, results in diploid populations being vulnerable to invasion by haploids. The rate of reversion cannot be too high for CBS138-derived diploids, because on the time scale of this experiment, diploid populations adapting in the no-FLC condition did not experience detectable haploid invasion. At 50 μg/mL FLC, the vulnerability to invasion is increased, perhaps because haploids with *PDR1* mutations have an even greater fitness advantage over diploids with *PDR1* mutations (Fig. 2C). At 100 μg/mL FLC, the additional FLC in the medium is enough to strongly select for mutations to *ERG11* or *ERG25* in addition to *PDR1*. Both of these genes are essential. Haploids and diploids thus appear to face different fitness landscapes, with diploids sometimes being able to out-adapt haploids, on the experimental timescale, by virtue of accessing heterozygous states that may be forbidden to haploids.

### Under increasing FLC concentrations, haploids and diploids adapt distinctly

The primary evolution experiment revealed greater divergence in adaptive paths between haploids and diploids at 100 μg/mL FLC than at 50 μg/mL, and the double-heterozygote (*PDR1 ERG11* or *PDR1 ERG25*) diploids adapted to 100 μg/mL FLC had higher MICs than evolved haploids (Fig. 2B). In light of these observations, we carried out a follow-up evolution experiment in which the concentration of FLC was dynamically adjusted upwards to 1024 μg/mL. In this experiment, we detected no shifts in ploidy. Notably, there were no shared genetic targets between haploids and diploids. Haploids adapted via mutations in *ERG3* and *OSH3*; diploids adapted by heterozygous mutations in *ERG25* plus multiple aneuploidies (Table 2). This experiment highlights the important role of aneuploidies in adaptation (discussed more fully below), and shows that there are situations in which haploids and diploids adapt via fully distinct paths.

The dependence of adaptive path on ploidy is at least partly driven by the interplay of dominance and gene essentiality. Haploid mutations to *ERG3* and *OSH3* were often loss-of-function; attaining a similar state in diploids would require two steps. Conversely, the heterozygous *ERG25* LOF mutations observed in diploids would be lethal in haploids, since *ERG25* is essential. Previous experimental evolution work comparing haploid and diploid yeast adaptation to azoles (e.g. [27], [43]) has not, to our knowledge, documented fully non-overlapping genetic targets, as our results show for *C. glabrata* in these conditions.

### K152E targets a recurrent fungal Erg11 resistance site

Erg11 (lanosterol 14α-demethylase) catalyzes a key step in ergosterol biosynthesis and is the target of azole drugs. *ERG11* missenses are common contributors to azole resistance in *C. albicans*, with a well-described hotspot at *Ca*K143 (=*Cg*K152) and surrounding residues [44], [45], [46], but *ERG11* mutations are only rarely present in clinical isolates of *C. glabrata* [47], [48], [49]. Engineered *C. glabrata* Erg11 substitutions corresponding to characterized *C. albicans* resistance mutations do confer azole resistance [50], and Ksiezopolska et al. observed recurrent *ERG11* mutations during evolution under fluconazole exposure, primarily in regimes combining fluconazole with anidulafungin or sequential exposure to both drugs, with K152 the most frequently altered Erg11 residue and K152E the most frequent substitution [51]. Thus, *ERG11* mutations are typically not a favored adaptive path for azole resistance in *C. glabrata*. Our observation of a high-fitness diploid clone carrying *ERG11* K152E/+ (in combination with *PDR1* V329F/+) is consistent with a heterozygote-advantage model, in which one wild-type allele preserves native Erg11 function while the K152E allele reduces fluconazole susceptibility. This inference is supported by the observation of an intermediate diploid-derived haploid lineage that, while dispensing with the second copy of all other chromosomes, maintained a chrE disomy that preserved the *ERG11* K152E/+ heterozygosity (Table 1).

### Heterozygous *ERG25* mutations and azole resistance

When Erg11 is inhibited by azole drugs, Erg25 catalyzes C-4 methyl oxidation of 14α-methylated sterol intermediates upstream of Erg3-dependent toxic diol formation [20]. Erg25 is essential under standard growth conditions in *C. glabrata* [52], so total inactivation of Erg25, which would sharply reduce flux through this alternative pathway, is not viable. In diploids, we observed one instance of heterozygous mutation in *ERG25* in the first evolution experiment, and four instances in the second. Of these five independent mutations, two are clearly loss-of-function, and two are missense substitutions at site D259 (D259H and D259Y), which has not been previously identified as a resistance locus. Relatedly, Zhou et al. (2024) recently identified recurrent heterozygous point mutations in *ERG251* during in vitro adaptation of *C. albicans* to FLC [53]. (In *C. albicans*, there are two paralogous C-4 sterol methyl oxidases, *ERG25* and *ERG251*.) They observed that these mutations confer azole tolerance but not classical resistance; resistance emerges when *ERG251* het-LOF is combined with trisomies of Chr3 and Chr6, which carry efflux pump genes including *CaCDR1* and *CaCDR2*. These observations are similar to ours: in the dynamic concentration experiment, heterozygous *ERG25* LOF mutations were always paired with trisomies that include chrF, which contains *CDR2* (also called *PDH1*). Thus, our data raise the possibility of a convergent adaptive pathway between *C. albicans* and diploid *C. glabrata* in which partial loss of C-4 sterol methyl oxidase activity, achieved by heterozygous mutation, is combined with an efflux-amplifying aneuploidy.

### Recurrent *ERG3* and *OSH3* co-mutation under high-FLC selection

As mentioned above, Erg3 catalyzes the final step in toxic diol production under Erg11 inhibition. While *ERG3* loss-of-function mutations alone confer azole resistance in *S. cerevisiae* [54] and *C. albicans* [55], in *C. glabrata* the picture is more nuanced. Geber et al. (1995) reported that *C. glabrata erg3*Δ deletion mutants are viable but not resistant to FLC [56]. Ksiezopolska et al. (2021) found that under selection for anidulafungin resistance, cross-resistance to FLC was associated with nonsynonymous *ERG3* mutation, with the missense D122Y conferring increased FLC resistance when tested by reintroduction, and reported that some strains that evolved *ERG3* frameshift or nonsense mutations retained wild-type susceptibility to FLC. Contrastingly, Gregor et al. (2023) reported increased FLC resistance in *C. glabrata ERG3* deletion strains, coupled with a growth rate defect [57].

Here, we have found that under selective pressure from very high levels of FLC, haploid *C. glabrata* repeatedly acquire concurrent *ERG3* and *OSH3* mutations. Among four independently evolved haploid clones, two carried an *ERG3* stop-loss and an *OSH3* missense; one carried an *ERG3* frameshift and an *OSH3* missense; and one carried an *ERG3* missense and an *OSH3* frameshift. *OSH3* (CAGL0J10780g) has not to our knowledge previously been reported as an acquired azole-resistance locus in *C. glabrata*. In *S. cerevisiae*, Ottilie et al. (2022) identified *OSH3* among genes mutated during in vitro evolution under posaconazole [58]. The oxysterol-binding proteins (Osh) are an evolutionarily conserved family of lipid-binding proteins, though Osh3 is unusual among Osh proteins in that it lacks sterol binding activity [59]. Osh3 deletion confers resistance to myriocin (ISP-1), a different class of drug, in *S. cerevisiae* [60]. We suspect that *OSH3* perturbation may be an important modifier of the sterol pool and/or membrane lipid organization resulting from *ERG3* dysfunction. We have separately observed that haploid *C. glabrata* colonies plated on YPD+FLC agar develop papillae that carry loss-of-function mutations in CAGL0J03916g (B.G.S., unpublished observations), currently annotated as the ortholog of *ScKES1/OSH4*, suggesting that multiple *OSH*-family genes can be selected as fluconazole-resistance modifiers in this organism. Further work will be required to elucidate the possibly synergistic mechanisms by which *ERG3* and *OSH3* mutations contribute to FLC resistance.

### Aneuploidy and adaptation to FLC

We found that adaptation by diploids was often accompanied by aneuploidy, in particular gains in the copy number of specific chromosomes. Natural (haploid) *C. glabrata* isolates can carry whole-chromosome and partial-chromosome aneuploidies, including whole-chrE and partial-chrG gains [61]. In experimental evolution, Ksiezopolska et al. (2021) found that selection under FLC enriched for chrE disomy in haploids. In our experiments, we observed one instance of chrE disomy in a diploid-derived haploid and one instance of partial trisomy in a diploid. We found, under two selection regimes involving FLC, that diploid *C. glabrata* recurrently acquired trisomies of chrC, chrF, chrG, and chrI, and these trisomies tended to co-occur. We did not detect aneuploidies in the no-FLC control populations. Possible drivers for these recurrent copy-number increases that have been implicated in azole resistance include the sterol regulatory transcription factor *UPC2A* on chrC [62]; the ABC transporter *CDR2/PDH1* on chrF [63]; the drug:H+ antiporter *CgQDR2* on chrG [64]; and the MFS transporter *TPO3* on chrI [65]. We observed some evidence that aneuploidies may provide a transient adaptive state: in one diploid replicate population adapting at 100 µg/mL FLC, we detected a *PDR1* V329F/+ *ERG11* K152E/+ diploid carrying chrC, chrF, and chrI trisomies at transfer 10. By transfer 20, we detected diploids with the same *PDR1* and *ERG11* mutations, but one retained only the chrC trisomy and the other had no trisomies.

### Limitations

The primary limitation of this study is that resistance mutations are inferred by parallelism in replicate populations, as well as concordance with prior published reports. Formal demonstration of the effect of particular mutations on both resistance and fitness remains a target for future work.

## Conclusions and further directions

We have shown here that spontaneous diploids are easily isolated from the *C. glabrata* type strain CBS138. Our results highlight that ploidy differences in *C. glabrata* can lead to different adaptive responses to the same selective pressure. At physiological concentrations of FLC, haploid and diploid *C. glabrata* share the well known *PDR1* adaptive pathway, but diploids additionally acquire heterozygous mutations to *ERG* genes that lead to genotypes that are both higher-resistance and higher-fitness. Diploids are also more likely to adapt to FLC via aneuploidy. At very high concentrations of FLC, diploid and haploid *C. glabrata* adapted through disjoint genetic mechanisms. Specific questions for further investigation include whether *ERG11* K152E/+ is a case of heterozygote advantage; how *ERG3* and *OSH3* mutations interact to lead to very high FLC resistance; why chrC, chrF, chrG, and chrI aneuploidies are repeatedly selected for under FLC; and whether these genotypes would be stably maintained in longer-term selection experiments. Reports of diploid clinical isolates of *C. glabrata* remain limited to date, but ploidy may not always be surveilled. In *C. auris* diploids are more virulent in a mouse model [66]; if and how diploidy affects virulence in *C. glabrata* remains a prospect for future work.

## Data availability statement

Supplementary File 1 contains all Supplemental Figures. Supplemental Files 2 and 3 are spreadsheets of all called mutations observed in the evolved clones, in the first and second evolution experiment, respectively. Sequencing data for this study is available at NCBI SRA <u>PRJNA1494094</u>. Strains described herein are available upon request.

