## Supplemental figures for "Diploidy alters the path of fluconazole adaptation in *Candida glabrata*"

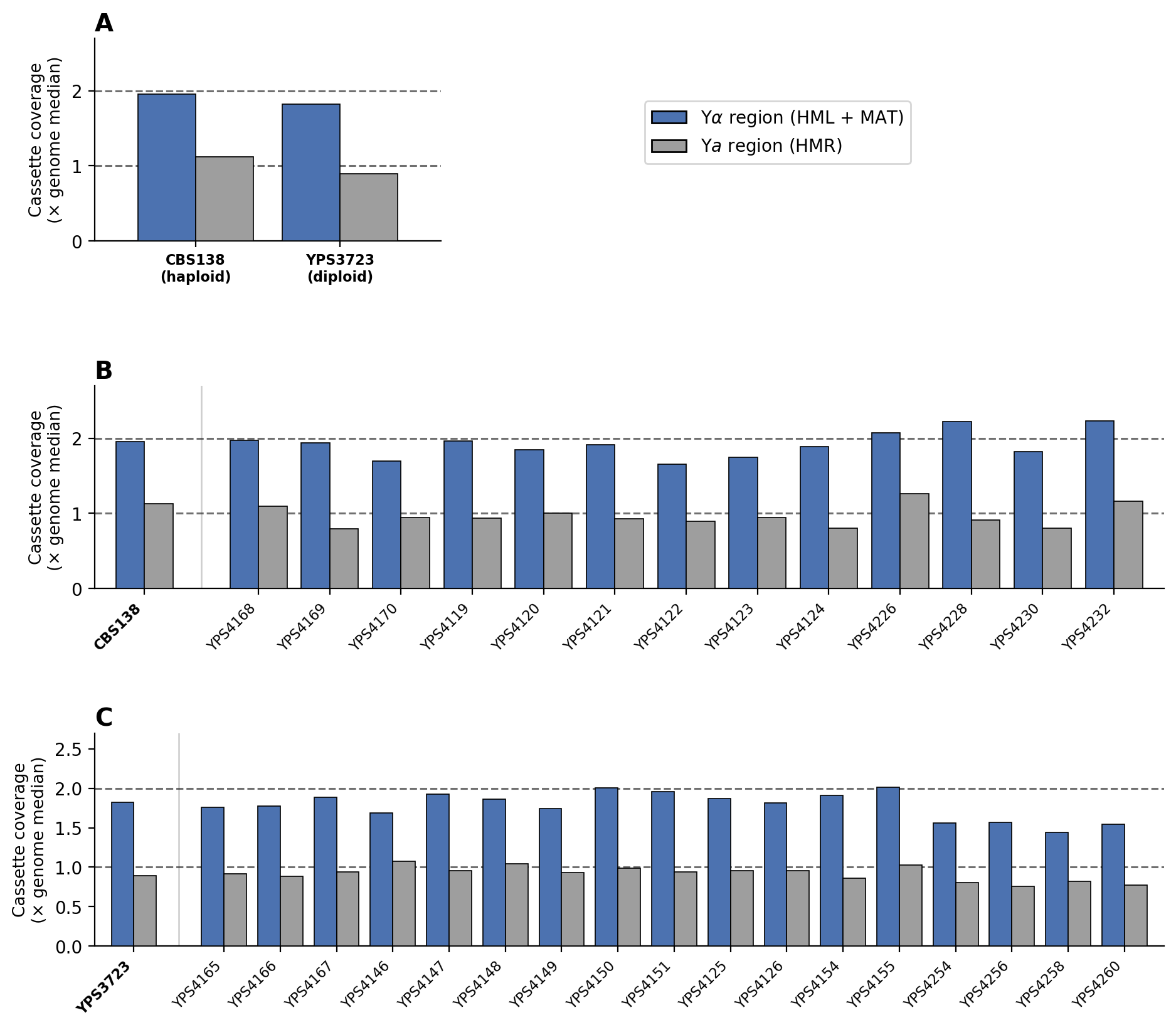

**Supplemental Figure 1.** The diploid ancestor YPS3723 (D-anc) is MATα/MATα and no evolution of mating type was observed. Bars show coverage of the Yα and Ya cassette regions, normalized to the genome-wide median. **A:** The ancestor CBS138 (H-anc) has the ~2:1 Yα:Ya ratio expected of a MATα haploid and the diploid ancestor YPS3723 has the ~2:1 Yα:Ya ratio expected of a MATα/MATα diploid; a MATa/MATα diploid would have a 1:1 ratio. **B:** No evidence of mating type evolution in clones evolved from the haploid ancestor. **C:** No evidence of mating type evolution in clones evolved from the diploid ancestor. Normalized coverage at mating type loci is depressed in YPS4254/56/58/60 because of multiple trisomies on other chromosomes.

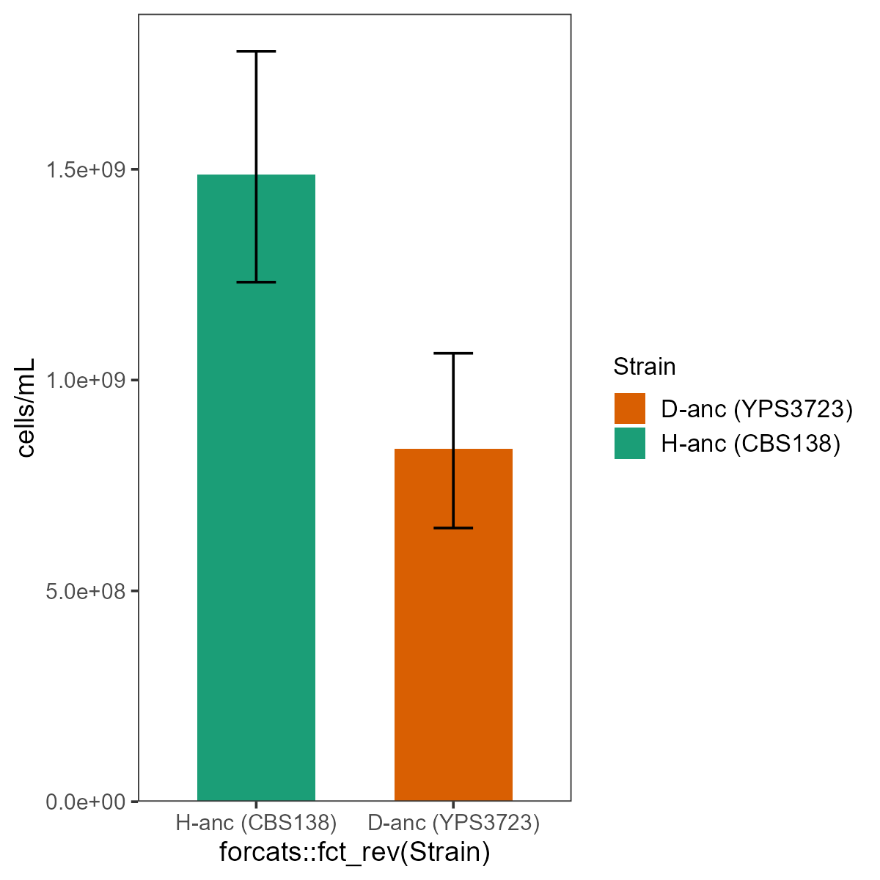

**Supplemental Figure 2.** Density of the two ancestors at saturation when grown in YPD, as measured by hemocytometer count.

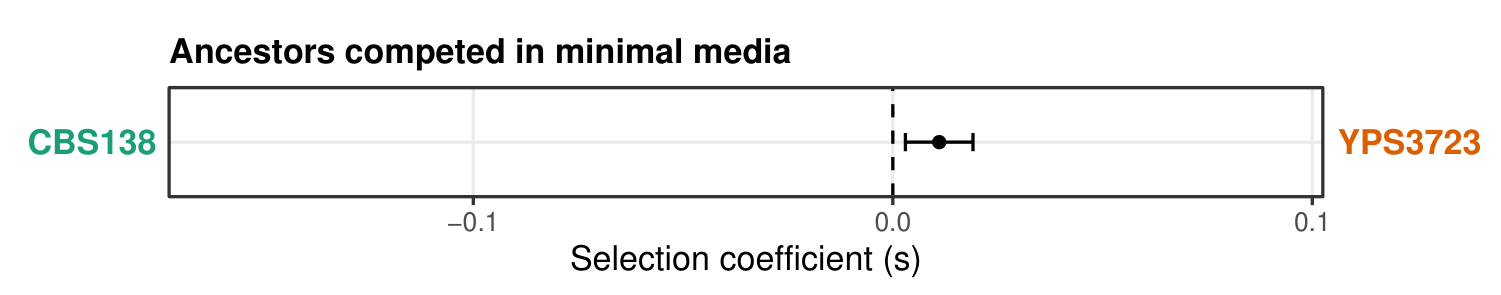

**Supplemental Figure 3.** A significant fitness difference between the diploid YPS3723 (D-anc) and the haploid CBS138 (H-anc) is not detected in minimal media (SD). The p-value for the mean *s* being nonzero is 0.07.

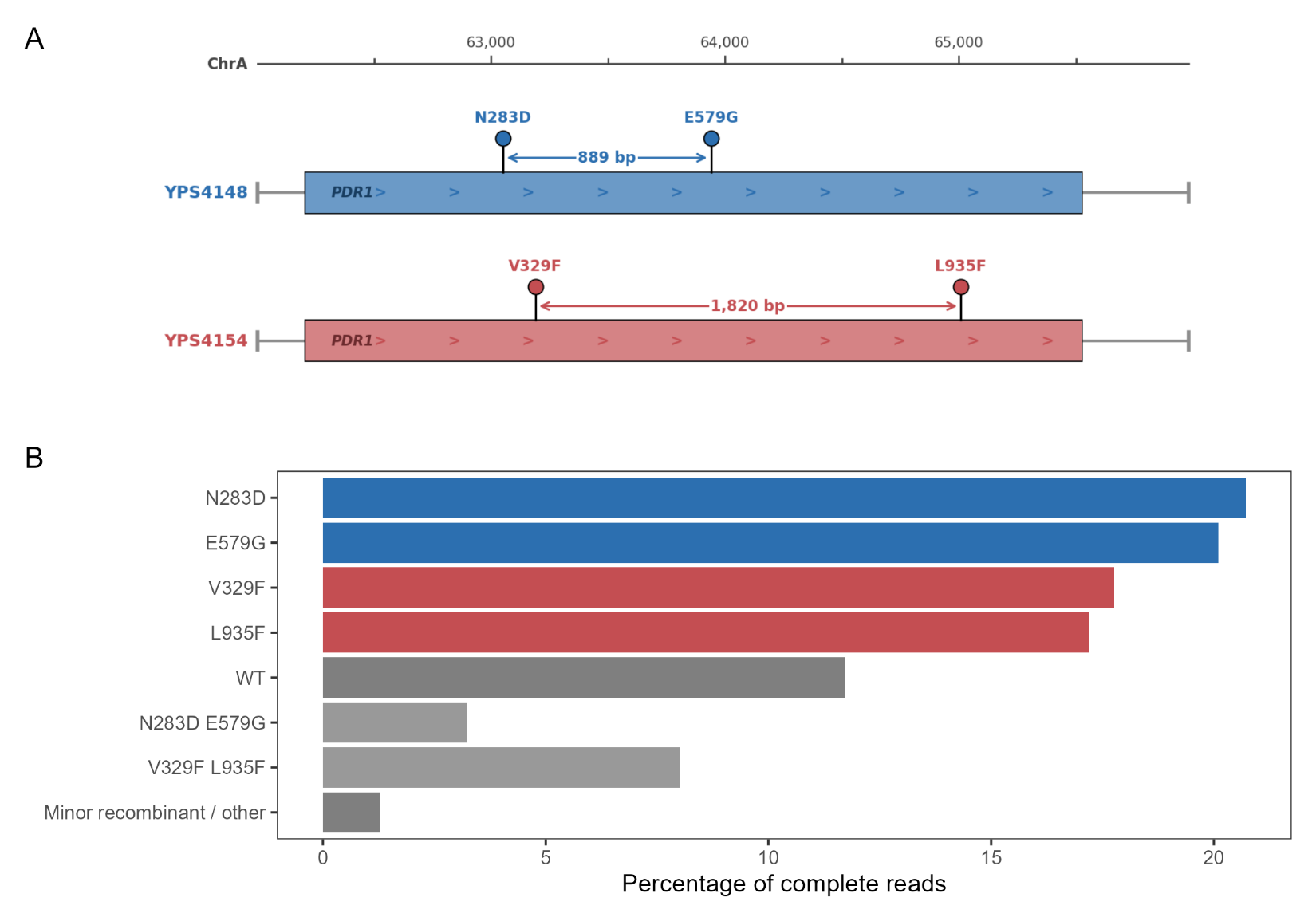

**Supplemental Figure 4.** The diploids heteroallelic for *PDR1* mutations have their mutations in *trans*. **A** is a schematic of *PDR1* and variant sites. PCR amplicons of *PDR1* from the diploids YPS4148 (D50-B1) and YPS4154 (D100-C1) were pooled for long-read sequencing. 1,887,895 reads spanned all four mutation sites, and the proportion of reads with zero, one, or two mutations is shown in **B**. The large majority of mutation-carrying reads have one mutation, implying that the heteroallelic *PDR1* diploids have their mutations on separate chromosomes. The increased proportion of *cis* reads for YPS4154 (D100-C1), compared to YPS4148 (D50-B1), is consistent with distance-dependent chimeric amplicons produced by PCR-mediated recombination; the zero-mutation (WT) reads are reciprocal products of the same process.
